# MoCoLo: a testing framework for motif co-localization

**DOI:** 10.1101/2024.01.04.574249

**Authors:** Qi Xu, Imee M.A. del Mundo, Maha Zewail-Foote, Brian T. Luke, Karen M. Vasquez, Jeanne Kowalski

## Abstract

Sequence-level data offers insights into biological processes through the interaction of two or more genomic features from the same or different molecular data types. Within motifs, this interaction is often explored via the co-occurrence of feature genomic tracks using fixed-segments or analytical tests that respectively require window size determination and risk of false positives from over-simplified models. Moreover, methods for robustly examining the co-localization of genomic features, and thereby understanding their spatial interaction, have been elusive. We present a new analytical method for examining feature interaction by introducing the notion of reciprocal co-occurrence, define statistics to estimate it, and hypotheses to test for it. Our approach leverages conditional motif co-occurrence events between features to infer their co-localization. Using reverse conditional probabilities and introducing a novel simulation approach that retains motif properties (e.g., length, guanine-content), our method further accounts for potential confounders in testing. As a proof-of-concept, MoCoLo confirmed the co-occurrence of histone markers in a breast cancer cell line. As a novel analysis, MoCoLo identified significant co-localization of oxidative DNA damage within non-B DNA forming regions that significantly differed between non-B DNA structures. Altogether, these findings demonstrate the potential utility of MoCoLo for testing spatial interactions between genomic features via their co-localization.

## Introduction

The increasing number of genomic datasets produced by high-throughput sequencing and prediction algorithms have revealed interactions between genomic features and biological processes(1–3). Although these interactions take many forms, their concept, derivation and evaluation remain embedded in the frequency of ‘co-occurrence’. Co-occurrence describes an event in which two or more features are present, which can be tested for their appearance together more often than would be expected by chance (4). On the other hand, ’co-localization’ refers to an event in which two or more features are both present in the same spatial region/proximity. While co-localization requires co-occurrence, the latter does not imply the former. Herein, we focus upon sequence motif interaction by introducing a criterion that requires the occurrence of a genomic feature within another feature and vice-versa. We refer to this criterion as reciprocal sequence co-occurrence and define metrics that enable characterization of co-localization using it.

Historically, for testing the co-occurrence of events two general approaches are used, one based on a Fisher’s exact test and another based on Monte-Carlo simulation (4,5). Statistical models rely on strict assumptions that may not always be suitable for genomic analyses. For example, parametric tests assume an *a priori* distribution that is oftentimes based upon independent events. These testing assumptions would be difficult to address since they involve finding the optimal model and parameters to characterize varying lengths of genomic regions that are often correlated between molecular features. While empirical methods may overcome strict modeling assumptions, they require simulations that take into account sequence properties (e.g., length, nucleotide content) to generate meaningful results. This type of sequence property-informed simulation often comes with the price of high computational costs and thus, may be difficult to achieve in the absence of an efficient algorithm.

Herein, we introduce MoCoLo (Motif Co-Localization) as a framework for direct testing of sequence-level co-localization using empirical methods coupled with a property-informed simulation algorithm. A class of hypotheses are constructed for testing the random occurrence of one feature in another feature and vice-versa (i.e., reciprocal occurrence). For hypothesis testing, a simulation method is introduced that incorporates sequence properties to ensure that the simulated data is representative of the properties embedded in the observed data such that differences in occurrence due to confounding factors are minimized. We demonstrate the method with two case applications for testing genome-wide co-localization between sequence-level molecular features of the same data type using histone modifications, and between different data types using alternative DNA (i.e., non-B DNA) structure-forming motifs (e.g., G-quadruplex DNA, Z-DNA, and mirror repeats) and 8-oxo-dG, an indicator of oxidative DNA damage.

## Results

### Overview of MoCoLo framework

MoCoLo is an approach to test for global, genome-wide reciprocal co-occurrence, i.e., co-localization. We describe our method within the context of two genomic features, feature 1 and feature 2 (F1, F2) (**Fig.1a**). Each defined by varying lengths and numbers of motifs (M1, M2). Interest is in addressing the question of whether these two feature motif libraries are co-localized and if so, to describe their co-localization by genomic region. This study provides a simulation-based approach to test co-localization of two genomic features, integrating the processes of hypothesis testing metric selection, property-informed simulation, and statistical evaluation.

**Fig.1.**
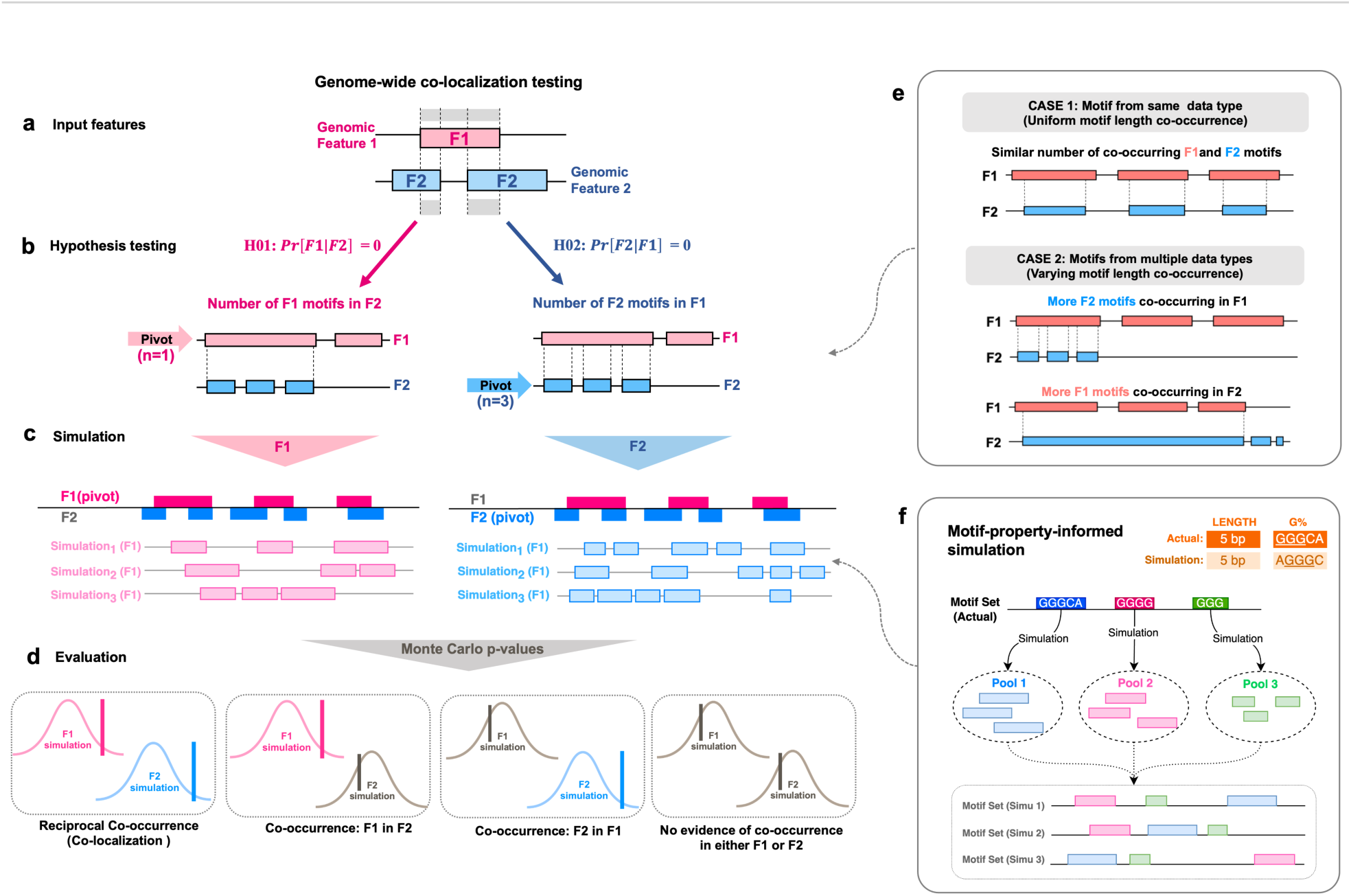
Overview of the MoCoLo framework for testing motif co-localizations. MoCoLo provides a simulation-based approach to test co-localization of two genomic features, integrating the processes of testing feature selection, property-informed simulation, and statistical evaluation. **(a) Input.** For testing co-localization, the input encompasses the genomic motif regions associated with features F1 and F2. **(b) Hypothesis testing.** A “pivot” feature is designated for hypothesis testing, recognizing that differences between the two motif data types can affect testing results (see also **e**). The co-localization assessment uses the number of the overlapping pivot features in the other as metrics. **(c) Simulation.** The motif-property-informed simulations will be performed in the next for each of the pivot motif group selected (see also **f**). It takes motif sequence characteristics into consideration to maintain the resemblance between the actual and the simulation groups. **(d) Significance evaluation.** MoCoLo determines the significance of co-localization by evaluate the two metrics reciprocally, incorporating Monte Carlo p-values in its results. If both hypothesis testing shows significant p-value, the two features are evaluated with “co-localization via reciprocal occurrence”. If only one side of tests shows significant p-value not the other, the two features have “co-occurrence of one in the other” but not co-localization. **(e) Motif type impact** on co-localization testing. Case 1 showcases co-localization when the length distributions of motifs from two features are alike, often originating from the same data type. Case 2 illustrates a co-localization scenario where motifs from the two features have contrasting sequence lengths. Here, a motif from one feature might overlap with several motifs from the other feature. The chosen testing hypothesis and simulation method in such situations can yield different results. **(f) Simulation design.** The design of the simulation method in MoCoLo emphasizes a motif-property-informed approach. This includes simulating individual motifs, constructing simulation pools, and assembling the simulated motif sets. Additionally, a “dynamic tolerance” is utilized to enhance computation efficiency and ensure a close resemblance between the actual and simulated data.

#### Reciprocal Co-localization Assessment

Our approach is designed for genome-wide reciprocal co-localization assessments (**Fig.1a**). Existing methods mostly test co-localization within the same genomic data type. While examining the notion of co-localization between motifs derived from different molecular data types, attention must be paid to the differences in sequence composition that define each data type (**Fig.1e**). It’s essential to consider the impact of difference in motif types on co-localization evaluation. In Case 1, similar motif length distributions, typically stemming from the same data type, might result in comparable counts of co-occurrence between two features (**Fig.1e, top**). Conversely, Case 2 depicts a situation where the motif lengths of the two features differ distinctly, potentially leading to one motif overlapping with multiple motifs from its counterpart (**Fig.1e, bottom**). Depending on the hypothesis and metric selected, these scenarios might produce varied results.

#### Duo hypotheses and testing metric

Therefore, we introduce two hypotheses that are both necessary to infer co-localization between F1 and F2 motif libraries (**Fig.1b,** “**Methods**”). The first hypothesis, H01, tests genome-wide, whether the number of F1 motifs in F2 motifs is greater than expected by random chance. Likewise, H02, tests genome-wide, whether the number of F2 motifs in F1 motifs is greater than chance. The two statistics for testing each hypothesis are based on estimates of conditional probabilities. A “pivot” feature needs to be designated for hypothesis testing, recognizing that differences between the two motif data types. The co-localization assessment uses the number of the overlapping pivot features in the other as metrics.

#### Sequence Property-Informed Simulation

As an empirical method, MoCoLo simulates expected data under a specified null hypothesis and compare it to the actual observed data (**Fig.1c**). It offers a simulation method informed by sequence properties to closely retain the characteristics of each motif groups. Unlike typical methods that utilize random re-positioning of regions, our method includes information on motif properties such as nucleotide composition in addition to motif length. The simulation method is developed by introducing new concepts such as simulation pool construction, motif sets assembling and dynamic tolerance, together to ensure a more nuanced simulation while maintain the computational efficiency (**Fig.1f**, “**Methods**”).

We applied MoCoLo to two case studies that focused on defining co-localization of different genomic and epigenomic features using same and different data type. In our first case study, we investigated the co-localization of two histone markers, H4K20me3 and H3K9me3. Case 1 provides a straightforward example of testing co-localization with direct length-only simulation and underscores the importance of two hypothesis tests, as a proof-of-concept. The second case study probed into the co-localization of non-B DNA-forming sequences with 8-oxo-dG lesion sites (different data type). We hypothesized that the distribution of 8-oxo-dG and non-B DNA-forming sequences within the genome differs between motif features. Case 2 highlights the need for feature-informed simulation in the testing framework. Here, both length and percentage of guanine (%G) of sequences were maintained to be similar and thus, minimize their differential effect in testing.

### The same-data-type co-localization testing of histone markers in breast cancer (Case1)

#### Background

Histone modifications play a significant role in regulating gene expression and maintaining genome stability. Among these modifications, H4K20me3 and H3K9me3 are well known for their roles in the formation of heterochromatin, a condensed form of chromosomal DNA associated with repression of gene expression. H4K20me3 plays roles in heterochromatin formation, gene expression repression (6) and genome stability regulation (7). Similarly, H3K9me3 is also crucial for heterochromatin formation (8,9). Our primary objective was to ascertain the extent of co-localization between H4K20me3 and H3K9me3 in the MCF-7 human breast cancer cell line utilizing the MoCoLo method as a proof-of-concept (**Fig.2a**).

**Fig.2.**
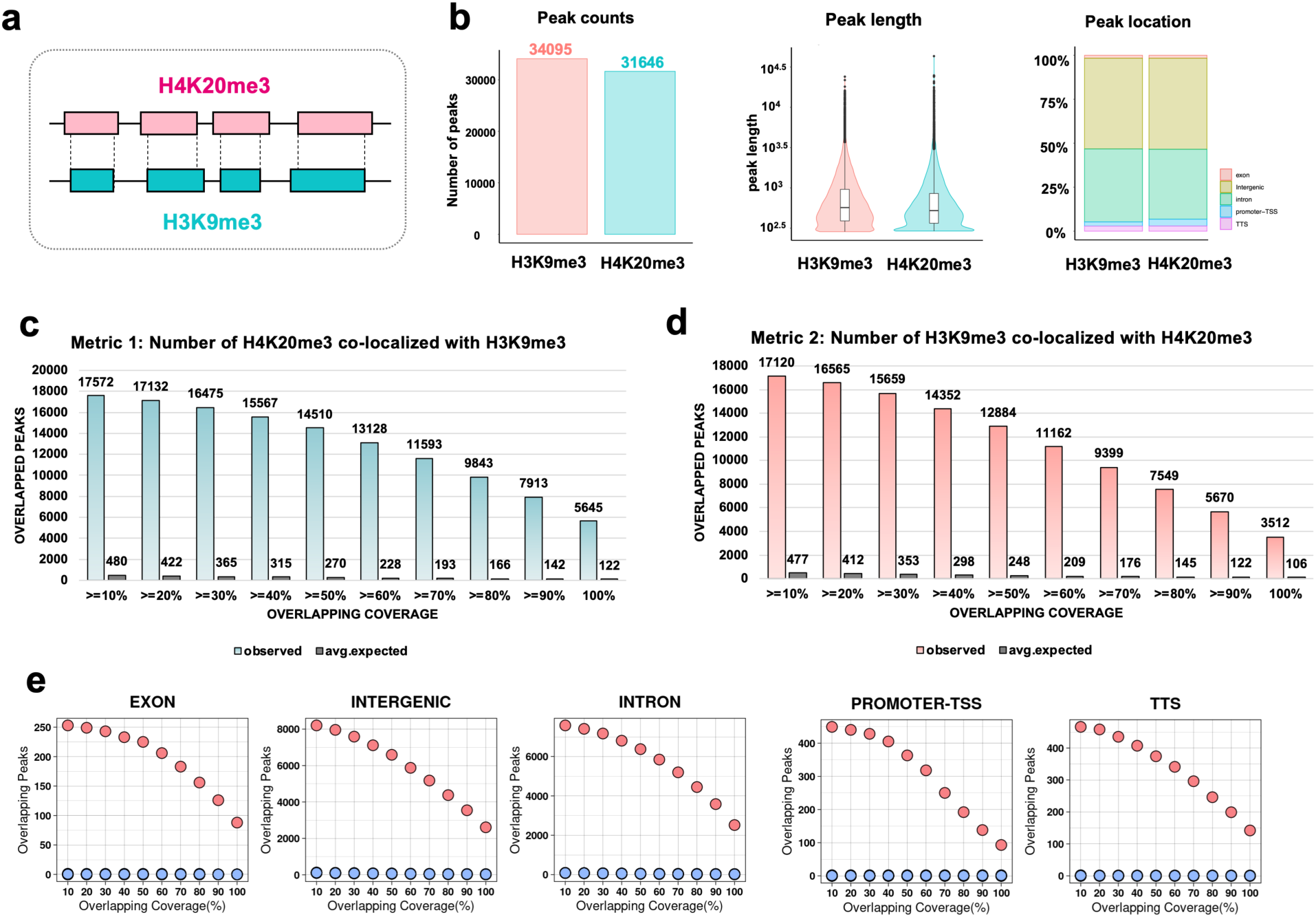
MoCoLo evaluates the co-localization of two histone markers, H4K20me3 and H3K9me3 (Case 1). **(a)** The objective is to assess the significance of co-localization between the H4K20me3 and H3K9me3 histone markers. **(b)** Peak details for the H4K20me3 and H3K9me3 markers in the MCF-7 breast cancer cell line. Both markers, from the same data type, display comparable peak length distributions: H4K20me3 has 31,646 peaks, and H3K9me3 has 34,095 peaks. (**c-d**) Genome-wide mapping utilizes H4K20me3 and H3K9me3 as pivots to evaluate two distinct metrics. The count of overlapping regions is assessed based on varying overlapping coverages (defined by the minimum intersection size). (**e**) Regional mapping examines the number of overlapping H4K20me3 peaks in co-localization across various genomic domains, such as exons, intergenic areas, introns, promoter-TSS, and TSS (red dot: observed; blue dot: expected).

#### Co-localization testing

H4K20me3 and H3K9me3 are both histone modification data generated from CHIP-seq experiments, thus sharing a data type and displaying comparable peak length distributions (**Fig.2b**). For our co-localization analysis, we conducted tests bi-directionally: one approach simulated H4K20me3 regions (n=31,646 regions) to establish the statistical distribution, and the alternate approach employed H3K9me3 regions (n=34,095 regions). Same lengths were retained while simulating histone peak regions (n=100). We then evaluated the test by using two metrics in term of the overlapped H4K20me3 and the overlapped H3K9me3. Both metrics showed significant differences in the observed group compared to the expected group, suggesting co-localization between these two histone markers. The count of overlapping regions is also assessed based on varying overlapping coverages (**Fig.2c-d**). In addition, we evaluated the co-localization at different genomic locations using the overlapped H4K20me3 as the evaluation metric. The results showed a higher number of overlapped regions in the observed group at exon, intergenic, intron, promoter-TSS (transcription start sites) and TTS (transcription termination sites) regions (**Fig.2e**).

The initial dataset for this case study underwent analysis via the segment annotation tool, ChromHMM. This tool delineates genomic regions by highlighting co-occurrence states between H4K20me3 and H3K9me3 (10). With MoCoLo we were able to formally test for co-localization between histone sites. Both approaches affirm the interaction between H4K20me3 and H3K9me3 sites, either in terms of co-occurrence using ChromHMM or co-localization using MoCoLo.

### The across-data-type co-localization testing of endogenous and exogeneous features of genomic instability (Case2)

#### Background

Genomic instability is a hallmark of cancer and other genetic diseases and can result from DNA damage from both exogenous and endogenous sources. Among the four DNA bases (A, T, C, G), guanine (G) has the lowest redox potential and thus has the highest propensity for oxidative damage (11–13). The oxidative lesion, 8-oxo-dG, therefore serves as a ubiquitous marker of oxidative stress (14,15) and is a pre-mutagenic lesion contributing to genome instability (11,16–18). Sequences that can adopt alternative (i.e., non-B) DNA structures are commonly enriched in guanines (11,19–21). Non-B DNA structures have also been shown to be co-localized with mutation hotspots in human cancer genomes (22,23) and can stimulate the formation of DNA double-strand breaks also jeopardizing genome stability (24–26). Further, 8-oxo-dG lesions have been shown to be enriched and/or refractory to repair in some types of non-B DNA (e.g., G4 and Z-DNA) (27–32), suggesting that these lesions may accumulate within such structure-forming sequences. The separate occurrence of 8-oxo-dG and non-B DNA-forming sequences are not uniformly distributed across the genome. The non-random distribution of 8-oxo-dG (27) may be due to increased oxidative damage potential and/or varied repair efficiencies within the local environment. We examined if the genome-wide co-localization of 8-oxo-dG and non-B DNA-forming regions and whether it differs between non-B types (**Fig.3a**), which include A-phased repeats (APR), G-quadruplex DNA (G DNA), Z-DNA (ZDNA), direct repeats (DR), inverted repeats (IR), mirror repeats (MR, also H-DNA), and short tandem repeats (STR).

**Fig.3.**
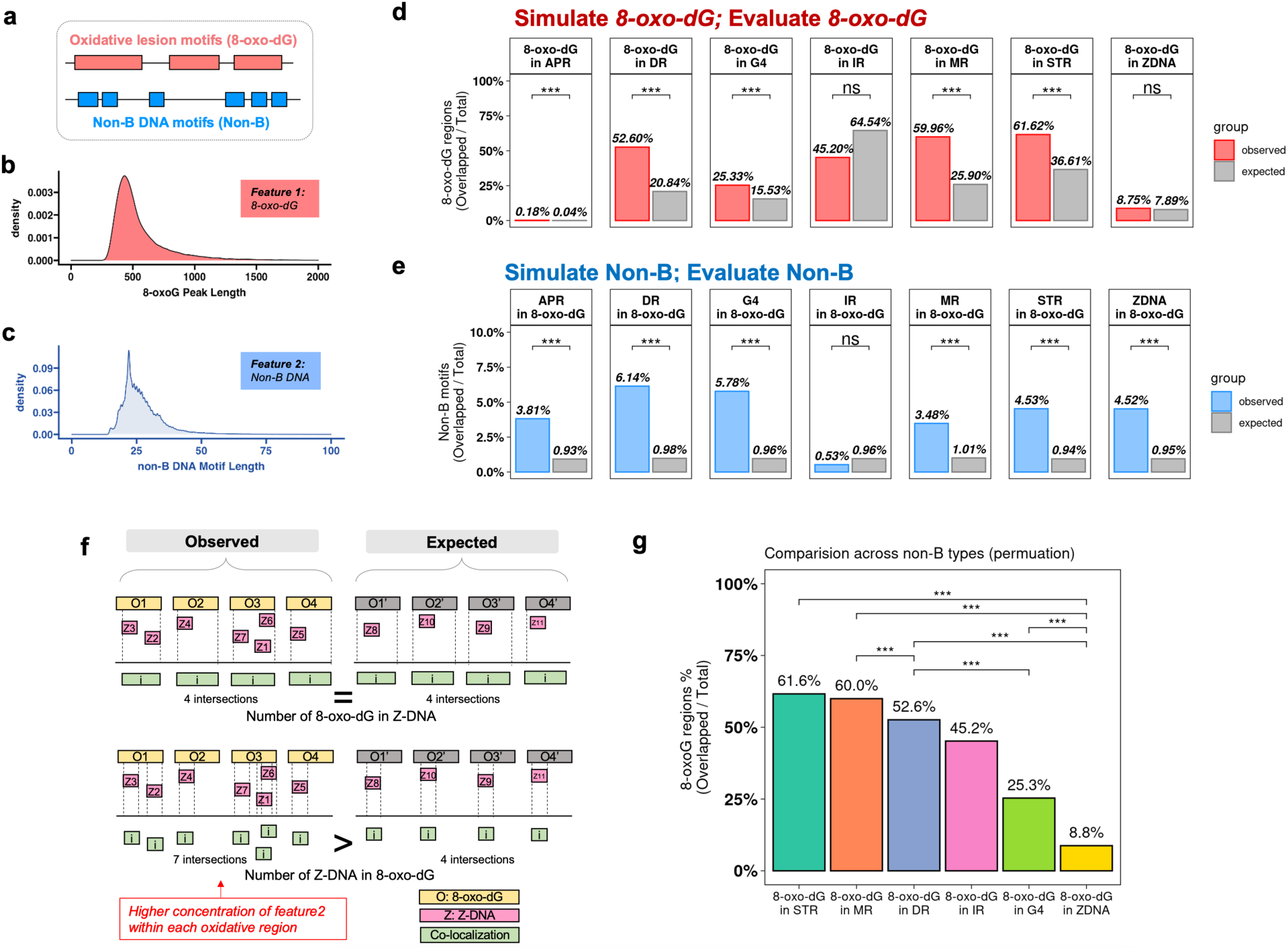
MoCoLo evaluates the co-localization of 8-oxo-dG and non-B DNA-forming regions (Case 2). (**a**) The overview of the genome-wide mapping of 8-oxo-dG peaks and non-B motifs. (**b-c**) The length distribution of 8-oxo-dG peaks (median, ∼500 bases) and non-B forming motifs (median, ∼25 bases). **(d)** The numbers of overlapped 8-oxo-dG regions (the observed) that co-localized with non-B DNA motifs by non-B type. 8-oxo-dG show significant co-localization with 6 non-B types except IR and Z-DNA. **(e)** The numbers of overlapped motifs of each non-B type that co-localized with 8-oxo-dG regions. Six non-B types show significant co-localization of their structure forming region and 8-oxo-dG region except IR. **(f)** While testing the co-localization between Z-DNA and 8-oxo-dG, there are significantly higher frequency of overlapped Z-DNA in the observed group while there is no significant difference of overlapped 8-oxo-dG. The explanation is that there is a high enrichment of Z-DNA in the certain 8-oxo-dG regions. Therefore, while counting ZDNA, there are higher overlapped Z-DNA (bottom) while the overlapped 8-oxo-dG regions stay the same (top). The observation highlights the need and benefits of using two-metric evaluation of co-localization and the importance of pivot feature selection. **(g)** Comparative analysis of co-localization between different non-B types and 8-oxoG. It investigates whether certain non-B types exhibit higher co-localization with 8-oxoG compared to others. The evaluation of co-localization by using the number of overlapped 8-oxoG regions as the metric and the testing result across non-B types.

#### Necessity of maintaining G-content in 8-oxo-dG region simulation

The accurate simulation of 8-oxo-G regions is intrinsically tied to preserving the G-content. When randomizing positions of 8-oxoG regions, it is imperative to retain the inherent G-content. This stems from the fundamental nature of the 8-oxoG motifs; by their very definition, they are expected to encompass a specific G-content. Omitting this essential characteristic would lead to a misrepresentation in the simulation. From this standpoint, it becomes evident that the preservation of G-content is an important for the simulation step in this case.

#### Testing results

The length of 8-oxo-dG regions from DIP-seq (**Fig.3b**) and the length of non-B motif (**Fig.3c**) show distinct difference. Notably, 8-oxo-dG peaks detected from DIP-seq experiments were overall large in length (median: ∼500 bases) as compared to non-B DNA motifs (median: ∼25 bases). This observation underscores the needs of reciprocal hypothesis testing (**Fig.1e**). Further, the sequence property-informed simulation method from MoCoLo was applied to 8-oxo-dG peaks (n= 50,027) for genomic region simulation (n=100) that retains guanine contents in addition to motif lengths.

We observed a significantly higher number of 8-oxo-dG regions co-localizing with five non-B DNA structures (MR, DR, STR, G4, and APR) in the observed group. Conversely, for IR and Z-DNA, the 8-oxo-dG regions did not exhibit significant co-localization when compared to other random genomic regions (**Fig.3d and Fig.S1a**). Furthermore, when evaluating using the non-B DNA motif count as the metric, we identified a significantly higher number of six types of non-B DNA-forming motifs that co-localized in 8-oxo-dG regions compared to the simulated group. These motifs include MR, DR, STR, G4, Z-DNA, and APR (**Fig.3e and Fig.S1b**).

The co-localization of APR-forming regions and 8-oxo-dG peak regions only indicate that APRs are located in proximity to the 8-oxo-dG region since A-tracts themselves do not contain guanines. This is because the 8-oxo-dG peaks from DIP-seq experiments are ∼500 bp while the A-phased repeats are ∼25 bp. Therefore, a 25-bp APR motif may co-localize within a 500-bp 8-oxo-dG region from DIP-seq peaks but does not mean that the one-base-specific oxidative guanine is located within the A-phased repeats themselves. The A-phased motifs are defined as three or more tracts of four to nine adenines or adenines followed by thymines, with centers separated by 11–12 nucleotides (33). The difference in peak sizes between the two data sets reflects a limitation of the current experimental technology to detect 8-oxo-dG within relatively smaller peak regions. It would be more fitting if the 8-oxodG sites can be detected in a narrower region or at single-base resolution.

### The dual hypothesis testing identified Z-DNA hotspots with 8-oxoG regions

Utilizing both “total overlapped 8-oxo-dG motifs” and “total overlapped non-B motifs” as evaluative metrics brings clarity to the intricacies of feature co-localization, as exemplified by the Z-DNA case. “Total overlapped 8-oxo-dG motifs” measures the total count of 8-oxo-dG regions that overlapped with non-B DNA, providing insights into the oxidative damage sustained by these motifs. In contrast, the “total overlapped non-B motif” captures the number of non-B DNA motifs present within 8-oxo-dG regions, signifying their placement within oxidatively damaged DNA regions.

For 8-oxo-dG regions that are overlapped with Z-DNA, the total number of 8-oxo-dG is not significantly higher in the observed group than random (**Fig.3d**). However, when we determined the total overlapped Z-DNA motifs within the 8-oxo-dG peak regions, the number is significantly higher in the observed group (p<0.001) than by random chance (**Fig.3e**). While these results may appear conflicting, it indicates a high number of overlapped Z-DNA-forming regions within each oxidative region and suggest that Z-DNA may be more frequently affected by oxidative pressures marked by 8-oxo-dG (**Fig.3f**).

### The post-testing comparison after co-localization testing

#### Analysis of comparing the co-localizations with 8-oxo-dG between various non-B types

MoCoLo provides further statistically testing functions to compare the co-localization of different non-B structure and 8-oxo-dG regions. The goal is to test the co-localization of across genomic features. In this case, example is the non-B DNA motif, which is stratified into 7 distinct types. This method is used to investigates whether a specific type of non-B motif demonstrates a more pronounced co-localization with the 8-oxo-dG feature than its counterparts.

To evaluate the co-localization between each pair of non-B types, we employ a permutation analysis (n=100). This involves reshuffling the non-B motif regions across the paired non-B types and conducting a subsequent co-localization analysis for each iteration to establish the null model. The count of overlapping 8-oxoG regions is utilized as the metric to compare co-localizations with oxidative regions across the seven non-B categories. These counts of overlapped regions are then normalized (by dividing by the total count of 8-oxo-dG regions or the respective non-B motif library sizes) to ensure comparability.

In terms of the overlapped 8-oxoG regions (**Fig.3g**), we observed significantly higher proportion of 8-oxo-dG regions were found to co-localize with MR (60.0%) than with DR (52.6%) and Z-DNA (8.8%). The co-localization of 8-oxo-dG and with STR (61.6%) and G4 (25.3%) are significantly higher than with the Z-DNA conformations. It also shows significantly higher frequency in DR than in G4 and Z-DNA.

The testing extension provides an alternative perspective to subgroups of genomic regions inherent to a singular genomic feature. Additionally, this approach melds both permutation (resampling within paired non-B types) and bootstrap (simulation of the 8-oxo-dG region) methodologies. This provide more insights in the co-localization and helps us understand how endogenous damage in the DNA and its structures are linked.

### Property-informed simulation ensures g-content retention in 8-oxo-dG simulations

#### Simulation design

A straightforward way to simulate genomic regions is to randomly place all regions independently While this satisfies length considerations, ensuring compositional accuracy, like matching nucleotide compositions, becomes challenging. The simulation here is not simply simulating the sequence. It is searching at genome wide with motif coordinates to find genomic regions whose sequences have the similar property to the actual motif (**Fig.4a**). Currently there is not a computation-effective workflow existing to simulate genomic regions with both length and g-content. To counter these inefficiencies, we introduced a new search strategy for simulation in MoCoLo (**Fig.1f**). Instead of a collective simulation of all motifs, motifs are simulated individually, populating a “simulation pool” tagged by motif traits such as length and composition. From this pool, we then select a motif set that mirrors our actual dataset. A built-in “dynamic tolerance” mechanism ensures efficient matching, preventing infinite loops by automatically adjusting the simulation tolerance, especially when an exact genome match is elusive.

**Fig.4.**
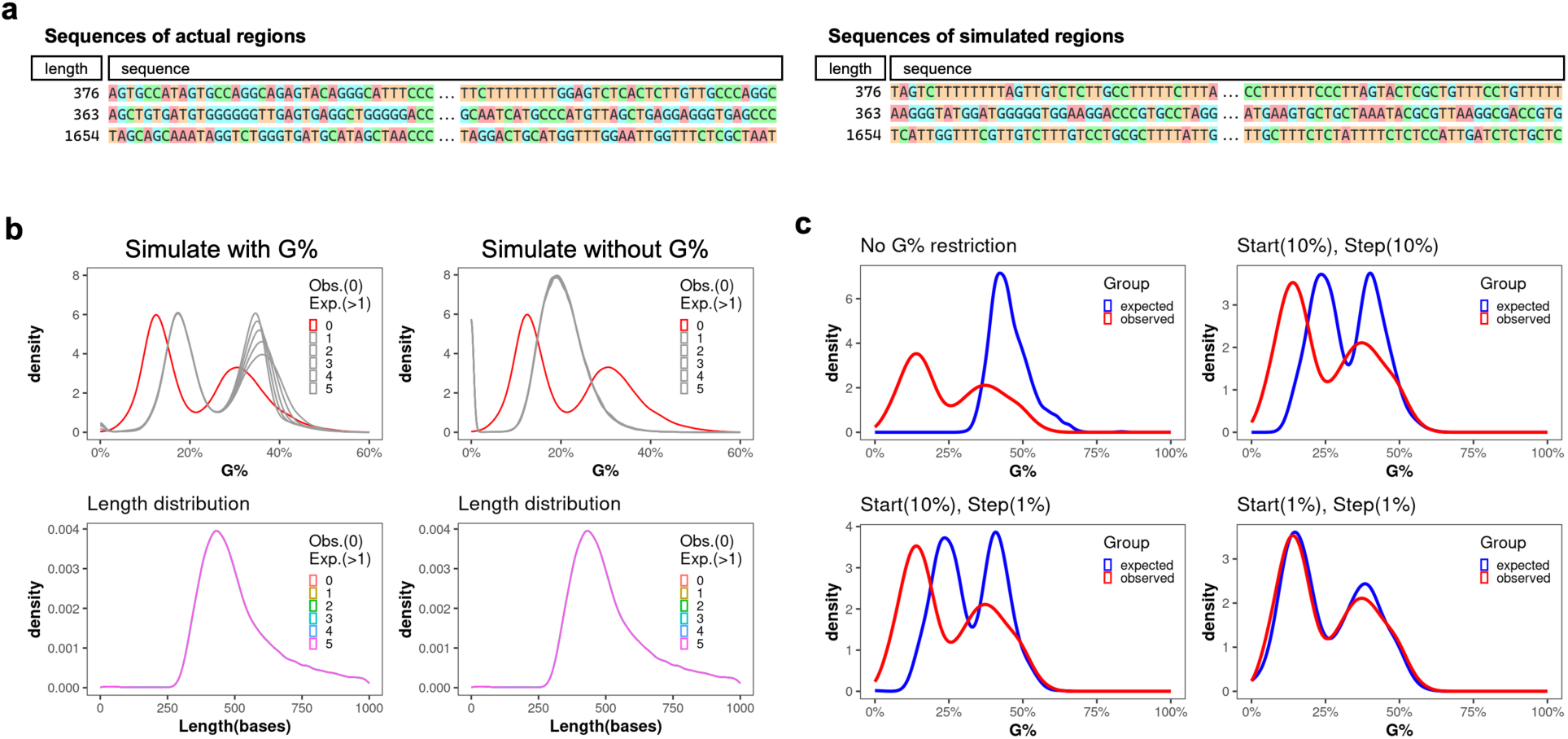
Property-informed simulation with dynamic tolerance maintains G-content of motif sequence. **(a)** The examples of property-informed simulation that retain the properties of motif sequence in terms of lengths and g-contents. **(b)** The distribution of G-Content of 8-oxo-dG region includes two G-content peaks for 8-oxo-G regions occur around 12.5% and 30.0%. G-content focused simulations underline the significance of G% for 8-oxo-dG. Overlooking G-content captures only length variation, whereas MoCoLo maintains both dual-peak G-content and length distribution, with a minor G-content shift hinting at the simulation’s tolerance. In the figure legend, 0 represent the actual data and 1-5 represent the simulation group. **(c)** The flexibility of the simulation is primarily influenced by two hyper-parameters: “starting tolerance (start)” and “incremental step (step).” The range for starting tolerance spans from zero — denoting an exact match to the G% of the original motif — to one, indicating no constraints on G%. If the starting tolerance is too stringent, the algorithm automatically adjusts the tolerance using defined increments set by the “incremental step.” The chosen values for “starting tolerance” and “incremental step” shape the attributes of the simulated groups, influencing their similarity to the real data. Top-left: An absence of G% constraint results in notable differences between simulated and actual groups; Bottom-right: Low start/step values result in heightened congruence between simulation and actual data, at the price of longer simulation time.

#### G-content variability

For 8-oxo-G regions, the G-content distribution presents two distinct peaks, approximately at 12.5% and 30.0%. A comparative analysis between simulations—with and without G-content restrictions—demonstrates the necessity of retain G% while simulating 8-oxo-dG regions. The property-informed simulation method in MoCoLo successfully preserves the dual-peak distribution, along with maintaining an identical length distribution (**Fig.4b, left**). In contrast, neglecting G-content in simulation retains only length distribution (**Fig.4b, right**).

#### Simulation parameters

The selection of parameters plays a pivotal role in simulation. We can observe a minor shift in the g-content distribution, which reflects the simulation tolerance (**Fig.4b, left-top**). Property-informed simulation in MoCoLo features “dynamic tolerance”. It is mainly regulated by two parameters: “starting tolerance (start)” and “incremental step (step).” Using the G% simulation as an example, the starting tolerance can vary from zero (0), indicating that the simulated motif should precisely reflect the G% of the actual motif, to one (1), which suggests no G% restrictions. In scenarios where the starting tolerance is excessively restrictive, the algorithm autonomously increases the tolerance in pre-defined increments determined by the “incremental step.” The specific values assigned to “starting tolerance” and “incremental step” dictate the characteristics of the simulated groups, subsequently affecting their resemblance to the actual data (**Fig.4c**). While using restrictive parameters ideally improves similarity, it might inversely affect computational efficiency, resulting in extended running time. Thus, users need to balance between efficiency and precision.

## Discussion

We introduce MoCoLo, a testing framework for genomic co-localization, which has several key innovations and advantages. First, MoCoLo employs a unique approach to co-localization testing that directly probes for genomic co-localization with duo-hypotheses testing. This means that MoCoLo can deliver more detailed and nuanced insights into the interplay between different genomic features. Second, MoCoLo features a novel method for informed genomic simulation, taking into account intrinsic sequence properties such as length and guanine-content. This simulation method enables us to identify genome-wide co-localization of 8-oxo-dG sites and non-B DNA forming region, providing a deeper understanding of the interactions between these genomic elements.

When applied to real-world data, MoCoLo revealed the significant co-localization of H4K20me3 and H3K9me3, vital for heterochromatin formation, in the MCF-7 breast cancer cell line. In addition, we were able to perform a genomic mapping between non-B DNA-forming regions and oxidatively damaged (8-oxo-dG) regions. Our results show significant co-localization of 5 types of non-B DNA-forming sequences within regions of 8-oxo-dG lesions. Our findings regarding G4 is also consistent with a previous report showing significant enrichment of potential G4 structures within 8-oxodG peaks compared to randomly distributed regions in the human genome, as predicted by sequence-based G4 models (34). In addition to the number of non-B DNA regions co-localized with 8-oxo-dG, we also calculated the total number of 8-oxo-dG regions co-localized with non-B. This additional metric revealed the high density of Z-DNA in 8-oxo-dG-containing regions. MoCoLo also provides capabilities to perform comparisons of co-localization status. As an example, we compared the co-localization status of the 7 non-B types with 8-oxo-dG and identify difference between these non-B types of their co-localization with oxidative regions. The 8-oxo-dG DIP-seq data was obtained from the MCF-10 breast cell line. Thus, it will also be informative to explore the same test in other cancer cell lines when the 8-oxo-dG data is available to perform comparisons.

There exist several strategies to indicate associations and co-occurrences in genomic studies (**Table 1**): **Monte-Carlo Based Approaches.** The design of MoCoLo relies on the principles of Monte-Carlo tests, which are non-parametric models that offer wide test statistics and randomization strategies. These tests, while affording flexibility, come with the inherent challenge of being computationally intensive, demanding precise customization. The degree to which data characteristics are preserved in a null model can significantly influence the conclusions drawn from Monte-Carlo simulations. In an endeavor to perfect these simulations, MoCoLo employs a property-informed simulation technique to uphold sequence properties. An innovative feature introduced is the “dynamic tolerance” in simulations, which modulates the tolerance level of sequence property differences between the observed and the simulated groups. The art of formulating a research question in Monte Carlo testing methods plays a pivotal role, as it directly corresponds to the chosen test statistic. A case in point would be the analysis of co-localization of two genomic features, F1 and F2. The query might revolve around whether F1 appears within F2 more than what random chance would suggest. Interestingly, such a proposition can also be viewed from an asymmetric perspective, mandating a diverse test statistic. In order to address both perspectives in a unified framework, MoCoLo introduces dual hypotheses for infer co-localization between F1 and F2 motifs and offers two distinct metrics to test each hypothesis.

**Table 1.**
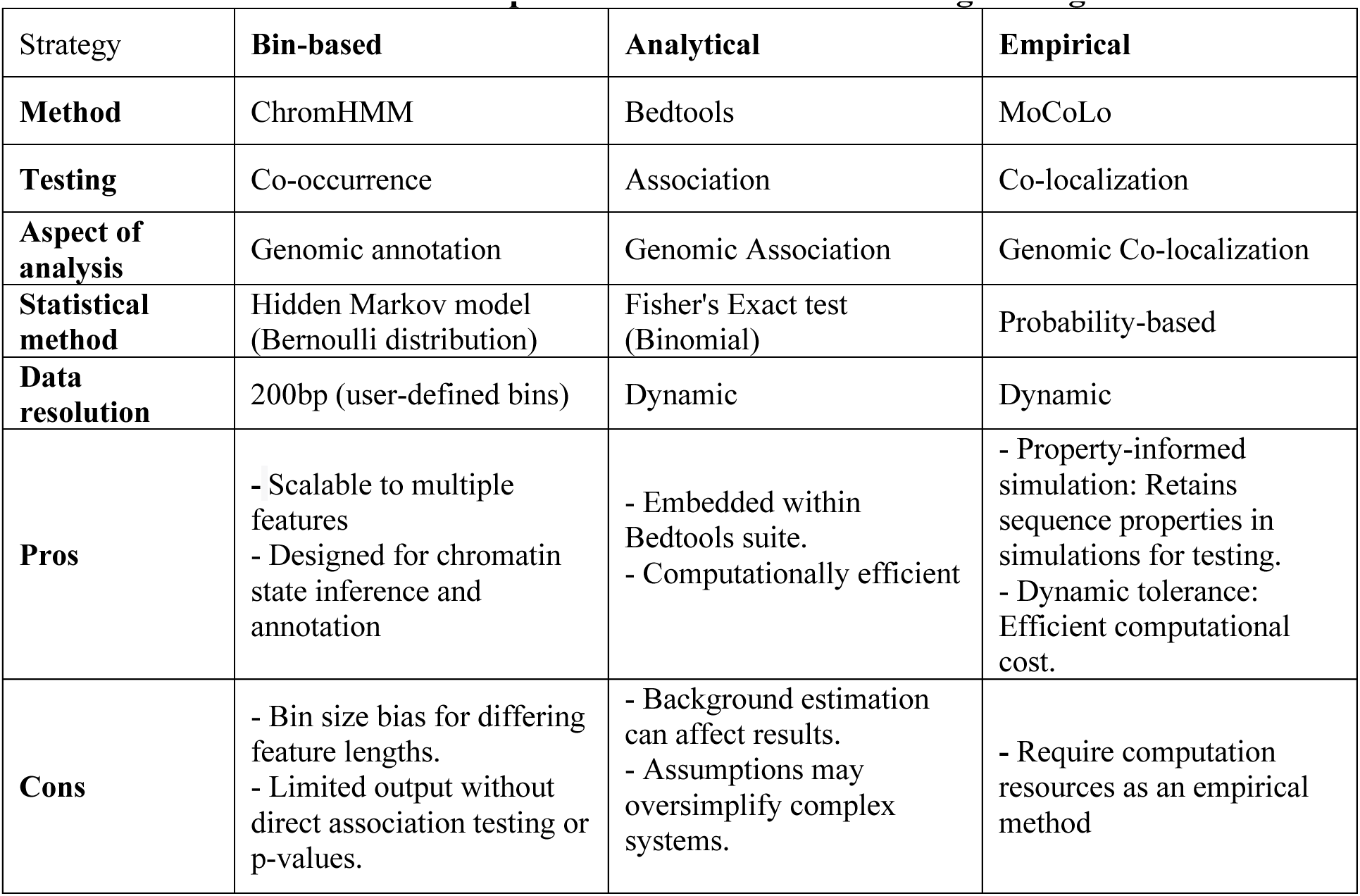
Overview of method comparison across different testing strategies.

| Strategy | Bin-based | Analytical | Empirical |
| --- | --- | --- | --- |
| **Method** | ChromHMM | Bedtools | MoCoLo |
| **Testing** | Co-occurrence | Association | Co-localization |
| **Aspect of analysis** | Genomic annotation | Genomic Association | Genomic Co-localization |
| **Statistical method** | Hidden Markov model (Bernoulli distribution) | Fisher's Exact test (Binomial) | Probability-based |
| **Data resolution** | 200bp (user-defined bins) | Dynamic | Dynamic |
| **Pros** | - Scalable to multiple features<br>- Designed for chromatin state inference and annotation | - Embedded within Bedtools suite.<br>- Computationally efficient | - Property-informed simulation: Retains sequence properties in simulations for testing.<br>- Dynamic tolerance: Efficient computational cost. |
| **Cons** | - Bin size bias for differing feature lengths.<br>- Limited output without direct association testing or p-values. | - Background estimation can affect results.<br>- Assumptions may oversimplify complex systems. | - Require computation resources as an empirical method |

### Approaches based on fixed-window segmentation

A prevalent approach in analyzing the co-occurrence of genomic elements involves segmenting them into multiple predefined window sizes, allowing for the calculation of statistics at the window level. Chromatin annotation tools such as ChromHMM, can be used to indicate the co-occurrence of two genomic features (the emission probability of a chromatin state). However, using a single fixed resolution during analysis may not be intuitive to decide resolutions especially when the two features in the testing has distinct length distribution. Therefore, despite the output (in terms of chromatin state annotations) of these tools can certainly be used as a foundation to study the co-localization of two genomic features, there are challenges existing such as 1) setting up bin-sizes, 2) restricted by statistical models, 3) no direct testing significant p-value provided in the output, as the primary objective of segmentation tools isn’t to test co-localization but to infer the co-occurrence in chromatin states.

### Analytical tests based on approaches

Basic analytical tests often rely on a straightforward null model, like that of Fisher’s exact test. When utilizing these tests, it’s crucial to assess if the data aligns with the null model and to understand the test’s resilience against any misalignments. Adopting an overly simplistic null model can lead to decreased P-values, heightening the chances of false positives. One implementation, Bedtools (35) provide implementation that can calculate the number of overlaps and the number of intervals unique to each feature. But it requires to infers the number that are not present in each feature as the universal background. Constructing the control set demands meticulous attention when using analytical tests rooted in a universe of regions. Any disparities between the case and control data sets in attributes such as genomic variability and aggregation could compromise the test’s assumptions, potentially resulting in false positives.

In summary, the main advantages of MoCoLo lie in its ability to handle dynamic and sequence-property-informed inputs, its reciprocal hypotheses testing, flexible simulation and its comprehensive output that allows for a more precise understanding of genomic feature co-localization.

## Materials and methods

### Testing hypotheses

We introduce two hypotheses that are both necessary to infer co-localization between F1 and F2 motif libraries. The first hypothesis, H01, tests genome-wide, whether the number of F1 motifs in F2 motifs is greater than zero. The second hypothesis, H02, tests genome-wide, whether the number of F2 motifs in F1 motifs is greater than zero. Formally, we introduce the following two hypotheses:

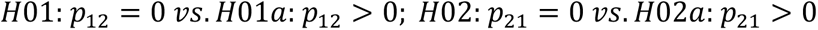

where:

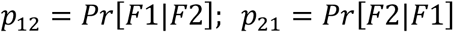

Below, we introduce two metrics for testing each hypothesis:

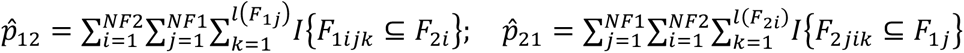

where *I*{·} is an indicator function, NF1 and NF2 are the number of motif libraries within features F1 and F2, respectively, and *l*(F_1j_) indicates the length of the j^th^ motif from F1 feature with *l*(F_2i_) the length of the i^th^ motif from F2 feature.

### Testing statistics

For gene-level overlap testing between two gene sets, denoted by G1 and G2, there exists options that are largely based on a Fisher exact test, with some popular choices being a Jaccard similarity coefficient and a hypergeometric distribution. If testing is two-sided, then we have no prior belief about direction and are simply testing whether the odds of success (‘overlap’) differs from 1 or not. On the other hand, one may be interested in a one-sided test of whether the odds of success (‘overlap of G1’) is greater (or less) in G2. In this context of a one-sided scenario, though not explicitly stated as such, one gene set is defined as fixed (i.e., ‘pivot’) that is compared against the other. We propose an analogous approach within a sequence context by introducing a feature variable pivot in which to conduct a (“two-sided”) test of association, the collection of which, H01: F1 in F2 and H02: F2 in F1 tests for co-localization association between features and the separation of which enables a ‘one-sided’ alternative. For pivot selection: we define “pivot selection” as the choice of reference feature to derive evaluation metrics. For testing H01, we quantify the total number of F1 motifs in F2, and thus, F2 is the pivot feature. Likewise, for testing H02, we quantify the total number of F2 motifs in F1, and thus, F1 is the pivot feature. Hence, we can evaluate co-localization by the reciprocal sequence co-occurrence by exchanging reference and query feature motifs.

### Property-informed simulation

Traditional brute force approaches simulate same-length genomic regions at random genome locations (35). This step fulfills the length requirement in simulation. However, the composition of the motif sequences in these simulated regions needed to be further checked and only those with similar nucleotide compositions (e.g., similar %G) are retained to fulfill the composition requirement. This can be computationally intensive and inefficient due to the potential non-existence of same-length regions with matching composition, which may lead to infinite loop situations.

To overcome these issues, we devised a novel optimal search strategy. As opposed to simultaneously simulating all motifs at once, instead, we simulated motifs individually and constructed a “simulation pool” that tags traits of interest for matching by motif length and composition. We then randomly sample a motif set (as set of simulated motifs with defined traits) from this pool that can be readily matched as the ‘random’ counterpart of the actual data motif set. Considering that another region with the exact same traits as the test region may not exist in the genome, with this approach, we were able to avoid the infinite loop created by enabling a “dynamic tolerance” that performs an automatic adjustment on the simulation tolerance.

### Data sources and processes

Histone Data. The ChIP-seq data of H4K20me3 and H3K9me3 in the human MCF-7 breast cancer cell line was downloaded from the NCBI Gene Expression Omnibus (GEO) under accession no. GSE143653 (36). The processed ChIP-seq data was download from GEO under the H4K20me3_BR_MCF7 (GSM4271383) and H3K9me3_BR_MCF7_rep2 (GSM4703869).

8-oxo-dG DIP-seq Data. The OxiDIP-Seq data was downloaded from the GEO database (GSE100234) (34). It contained the genome-wide distribution of 8-oxo-dG accumulation in human non-tumorigenic epithelial breast cells from the MFC10A human cell line. The processed peaks data were provided by the author in bed format.

Non-B DNA forming motifs. Non-B DNA-forming motifs were extracted from the updated version Non-B DB v2.0 database (human hg19 reference genome) (33). An update to correct the A-Phased repeat motifs data was received from Frederick National Laboratory for Cancer Research. It includes 13,966,212 motifs covering seven types of non-B structures: A-phased repeats (APR), G-quadruplex DNA (G DNA), Z-DNA (ZDNA), direct repeats (DR), inverted repeats (IR), mirror repeats (MR, also H-DNA), and short tandem repeats (STR).

### Function implementation

The functions bedtools_shuffle() and bedtools_random() from the ‘valr’ package are utilized to sample genomic regions at genome-wide. The “within” paratmeter is used to control whether to perform the with-in chromosome simulation or not. The bedtools_coverage() is utilized to quantify the overlapped regions between motifs from two genomic regions. Only with the length of overlapped region greater than 0 are the two regions considered co-localized. The visualization functions are implemented with the ‘ggpplot2’ package as well as the ‘ComplexHeatmap’ package. The significance annotation function in the visualization is from the ‘ggpubr’ package.

### Statistical Significance

For the evaluation of statistical significance in the co-localization testing, a Monte-Carlo based p-value is computed. This is executed for each formulated hypothesis. The computation involves a systematic comparison between metrics derived from both simulated and observed datasets. Specifically, the assessment quantifies the proportion wherein the metrics extracted from the simulated datasets surpass the corresponding metrics derived from the actual observed datasets.

## Key Points

□ MoCoLo framework provides a novel method for analyzing spatial interactions of genomic features at sequence-level using reciprocal co-occurrence.
□ Property-informed simulation in MoCoLo minimizes confounding factors, enabling robust genomewide co-localization assessments.
□ Through case studies, MoCoLo demonstrated its utility in unveiling significant co-localizations, aiding in deeper molecular understanding.

## Contributions

Conceptualization JK, QX, KMV, MZF, IDM; Methodology and Formal Analysis, JK, QX, BTL; Writing – Original Draft Preparation, JK, QX; Editing of Original Draft, KMV, MZF, IDM; Funding Acquisition, JK, KMV

## Availability of data and materials

The MoCoLo package is available under a GPL-3.0 license in the Kowalski Lab GitHub repository. The supporting script for the case study analyses for the paper can also be accessed in the scripts folder in the same repository(https://github.com/kmlabdms/MoCoLo).

